# The challenge of sequencing *Chlamydia trachomatis* and other bacterial sexually transmitted infections genomes directly from clinical swabs: the optimum solution

**DOI:** 10.1101/2024.11.23.624631

**Authors:** Karina Büttner, Vera Bregy, Fanny Wegner, Srinithi Purushothaman, Frank Imkamp, Tim Roloff Handschin, Mirja H Puolakkainen, Eija Hiltunen-Back, Dominique Braun, Ibrahim Kisakesen, Andreas Schreiber, Andrea Carolina Entrocassi, María Lucía Gallo Vaulet, Deysi López Aquino, Laura Svidler López, Luciana La Rosa, Adrian Egli, Marcelo Rodriguez Fermepin, Helena MB Seth-Smith, the ESCMID Study Group for Mycoplasma, Chlamydia infections (ESGMAC)

## Abstract

Rates of bacterial sexually transmitted infections (STIs) are rising and accessing their genomes provides information on strain evolution, circulating strains, and encoded antimicrobial resistance (AMR). Notable pathogens include *Chlamydia trachomatis* (CT), *Neisseria gonorrhoeae* (NG) and *Treponema pallidum* (TP), globally the most common bacterial STIs. *Mycoplasma genitalium* (MG) is also a bacterial STI which is of concern due to AMR development. These bacteria are also fastidious or hard to culture, and standard sampling methods lyse bacteria, completely preventing pathogen culture. Clinical samples contain large amounts of human and other microbiota DNA. These factors hinder the sequencing of bacterial STI genomes. We aimed to overcome these challenges in obtaining whole genome sequences, and evaluated four approaches using clinical samples from Argentina (39), Switzerland (14), and cultured samples from Finland (2) and Argentina (1). First, direct genome sequencing from swab samples was attempted through Illumina deep metagenomic sequencing, showing extremely low levels of target DNA, with under 0.01% of the sequenced reads being from the target pathogens. Second, host DNA depletion followed by Illumina sequencing was not found to produce enrichment in these very low load samples. Third, we tried a selective long-read approach with the new adaptive sequencing from Oxford Nanopore Technologies (ONT), which also did not improve enrichment sufficiently to provide genomic information. Finally, target enrichment using a novel pan-genome set of custom SureSelect probes targeting CT, NG, TP, and MG followed by Illumina sequencing was successful. We produced whole genomes from 64% of CT positive samples; from 36% of NG positive samples, and from 60% of TP positive samples. Additionally, we enriched MG DNA to gain partial genomes from 60% of samples. This is the first publication to date to utilize a pan-genome STI panel in target enrichment. Target enrichment, though costly, proved essential for obtaining genomic data from clinical samples. This data can be utilized to examine circulating strains, genotypic resistance, and guide public health strategies.

**Graphical Abstract:** 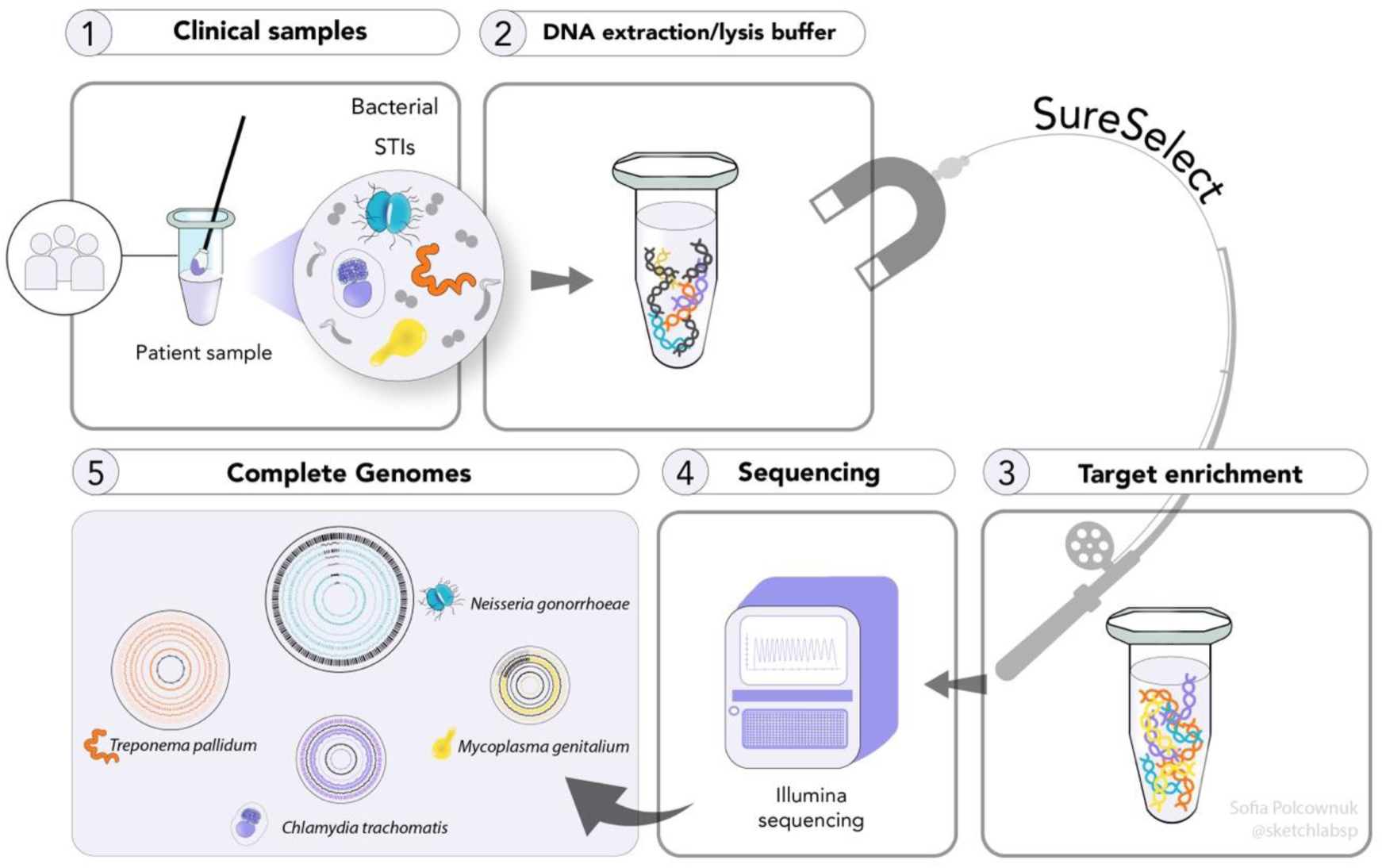

**Impact statement:** Genome data on circulating sexually transmitted infections (STIs) is important to better understand transmission networks, antimicrobial resistance and to guide treatment decisions. For many bacterial STIs, this information is difficult to obtain, as the bacteria are fastidious, in some cases intracellular, and often recalcitrant to culture. We have developed and tested a target enrichment STI panel of baits to capture whole genomes of *Chlamydia trachomatis*, *Neisseria gonorrhoeae*, *Treponema pallidum,* and *Mycoplasma genitalium* with approximately 50% success in genome sequencing for the first three pathogens. We compare this against other sequencing and enrichment methods, which did not provide sufficient data for genome analysis. This panel approach shows potential for clinical samples carrying these pathogens and can potentially also be developed for further pathogen groups.

**Data summary:** All illumina sequence data, with human read data removed using Hostile (1) and KrakenTools (https://github.com/jenniferlu717/KrakenTools), is deposited with the European Nucleotide Archive (ENA) under project number PRJEB72167.

## Introduction

The global burden of bacterial sexually transmitted infections (STIs) recently reached 374 million new cases annually (2) after increasing in recent years. Between 2010 and 2019, rates of chlamydia and syphilis increased, while gonorrhoea remained high, according to the Global Burden of Disease study (3,4), continuing a trend in rising STI numbers that has been seen over decades. Asymptomatic infections are frequent and can result in unknowing transmissions of STIs. In females STIs may result in pelvic inflammatory disease (PID) and scarring of the upper reproductive tract which is a risk factor for infertility and extrauterine pregnancy. In addition, antimicrobial resistance (AMR) is an important concern for some STI pathogens, with the prospect of untreatable gonorrhoea infection (5,6).

Chlamydia (caused by the bacterium *Chlamydia trachomatis*, CT) is globally the most common bacterial STI, with 129 million estimated new infections in 2020 (7), in comparison to 82 million new infections with gonorrhoea (caused by the bacterium *Neisseria gonorrhoeae*, NG) (8), which is increasingly resistant to many antimicrobial treatments leading to multi-drug resistant NG strains being transmitted globally (5). Syphilis (caused by the spirochaete bacterium *Treponema pallidum* subsp. *pallidum*, TP) is a third notifiable disease and is increasingly found to be macrolide resistant, although is susceptible to penicillin. *Mycoplasma genitalium* (MG) is commonly identified in samples diagnosed with STI diagnostic panels. MG rapidly develops AMR against macrolides and is a relevant pathogen in the setting of recurrent urethritis. These four pathogens are difficult to grow in the laboratory: CT is an intracellular bacterium, only able to grow within human host cells (9); NG is a fragile bacterium, requiring rapid processing and specific media; TP is exceedingly difficult to grow under standard laboratory conditions; and MG is fastidious and extremely slow to grow in culture (10). These STIs often occur in combination and the clinical signs and symptoms if present may be similar and hard to distinguish. Due to these diagnostic challenges new approaches are necessary.

Treatment of STIs requires accurate diagnosis, and rapid determination of underlying AMR is crucial for efficient treatment and the management of STI syndromes. For public health purposes, information is required on the circulating strains, patterns of transmission, and the extent of AMR. Genomic analysis allows the highest typing resolution, enabling comparison of bacterial strains at the level of every single nucleotide in the whole genome. With this data, strain relatedness can be delineated, and resistance and virulence determinants identified.

Culture free techniques for genome analysis of CT exist, with several possibilities for whole genome sequences (WGS) generation (11). Some of these techniques rely on having intact bacteria in the clinical samples. This is often impossible given current molecular diagnostic techniques, where samples go straight into lysis buffer for stabilisation of nucleic acids. These methods make culture of the species impossible, which is also a problem for AMR phenotyping, which is particularly relevant for NG, TP and MG (12). The most successful technique to date is that of target enrichment, also known as hybrid capture and sequence capture. Target enrichment, in various forms, has been used to generate whole genomes from bacteria in many studies (13–21) particularly for CT (22–28), NG (29) and TP (30–35) as well as other use-cases including organisms, RNA capture, specific targets (36–41), and one panel test including whole bacterial genomes involved in tick-borne diseases (42).

We aimed to explore techniques to gain bacterial STI WGS data from clinical samples, using deep sequencing, host DNA depletion, Oxford Nanopore Technologies (ONT) adaptive sampling, and we show the first data using a pan-genome target enrichment panel across bacterial STIs.

## Methods

### Clinical sample collection and diagnosis

At the University of Buenos Aires (UBA), rectal swabs from patients with symptoms compatible with proctitis were collected between September 2017 and August 2023. Adult patients (18 years of age and older) were prospectively included from a public hospital (Hospital Juan A. Fernández) and a private health centre (Centro Privado de Cirugía y Coloproctología) in Buenos Aires, Argentina. Patients had anorectal symptoms (anal discharge, rectal bleeding, rectal tenesmus, and proctalgia) and sexual risk factors suggestive of STIs. Patients were treated according to current STI guidelines (43,44). Rectal swabs were placed in 1.5 ml of 2-SP medium supplemented with 2% Foetal Calf Serum (FCS) and antibiotics (gentamicin 25 μg/mL, vancomycin 0.5 mg/mL, and amphotericin 2.5 μg/mL) for *C. trachomatis* culture and stored at -80°C.

From the University Hospital Zurich (USZ), all samples were from men who have sex with men (MSM) living with HIV, who were in the main asymptomatic, provided as pooled (urethral, anorectal, pharyngeal) swab samples using cobas PCR Media Uni Swab Sample Kit (Roche) deriving from a systematic testing strategy within two HIV cohorts (45,46).

Finnish samples from the Helsinki University Central Hospital (HUCH) were from MSM attending the outpatient sexually transmitted infection (STI) clinic because of symptoms suggestive of LGV or because notified by an infected partner were tested for *Chlamydia trachomatis* and *Neisseria gonorrhoeae*. From each patient, one rectal swab was transported in Aptima Swab Transport Medium (Hologic) for *C. trachomatis* and *N. gonorrhoeae* testing at the diagnostic laboratory of the Helsinki University Central Hospital (Aptima Combo 2 Assay, Hologic). A second rectal swab from each patient was placed in Universal Transport Medium (UTM, Copan) for *C. trachomatis* culture and stored at -70°C.

At the IMM, DNA extraction from USZ clinical samples was performed on a QIASymphony SP/AS instrument using the QIA DSP Virus/Pathogen Kit (Qiagen). We performed an in-house developed and accredited multiplex-PCR diagnostic real time (RT)-PCR panel based on Lightmix Kits from Tib Molbiol on all USZ clinical samples (47), and also retrospectively on all UBA samples to ensure data uniformity.

### Culture of Chlamydia trachomatis, DNA extraction and storage / transport

One cultured sample from UBA was used as a spike in the ONT adaptive sampling experiment. At UBA, LLC-MK2 cells (epithelial kidney cells from Rhesus Monkey ATCC CCL-7) were maintained in 10% fetal calf serum (FCS, Internegocios, FBI) supplemented minimal Eagle’s medium (MEM, Gibco) with 2 mM glutamine (Gibco), 1.5 g/L sodium bicarbonate, 1 mM non-essential amino acids (Gibco), and 50 μg/mL gentamicin and grown at 37°C under 5% CO_2_ (48). The infection medium for the culture of CT, in 25cm^2^ culture bottles comprised of this plus glucose 56 mM, cycloheximide 10 μg/mL, gentamicin 0.05 mg/mL, vancomycin 0.5 mg/mL, and amphotericin 2.5 μg/mL. For infection determination, duplicate samples were grown in shell vials, and cells on cover slips were fixed with pure methanol. FITC labelled antibodies against CT lipopolysaccharide (LPS) were used for staining (Goat X *Chlamydia trachomatis* AB1120F, Millipore). Samples were sub-cultured up to three times where necessary, and samples showing CT growth were stored in 50% FCS at -80°C. Infected LLC-MK2 cells were harvested, and DNA was extracted using the Quick-DNA MiniPrep kit (Zymo Research Corporation, Irvine, USA). The *ompA* gene was amplified using a nested PCR technique (49) and Sanger sequencing (Macrogen, Inc) (50).

Two cultured samples from HUCH were used as spikes in the ONT adaptive sampling experiment. At HUCH, the rectal swabs were cultured in McCoy cells (originally obtained from Flow Laboratories Limited, Irvine Scotland)(51). Cells in 24-well plates (some with glass coverslips) were inoculated with UTM and centrifuged at 3000× g at 30°C for 1 h (52). BHK-21 medium (based on Glasgow minimal essential medium, Sigma Aldrich) supplemented with 10% fetal calf serum, 2 mM glutamine, 20 μg/mL gentamicin, 50 U/mL nystatin, 100 μg/mL vancomycin, and 0.5 μg/mL cycloheximide was added, and the plates were incubated in 5% CO2 at 35°C for 48 h. The growth of *C. trachomatis* was detected by direct immunofluorescence staining of the coverslips with a Pathfinder *Chlamydia* Culture Confirmation System (Bio-Rad, Hercules, CA, USA). The infected cells from wells with no coverslips were collected in sucrose–phosphate–glutamate, pH 7.2 (SPG), and slowly frozen to −70°C. To increase the amount of bacteria, a passage was done (the cells were re-inoculated into uninfected McCoy cells with the procedure described above). Infected McCoy cells were harvested and DNA was extracted with Maxwell RSC Cultured Cells kit (Promega).

### Pathogen quantification and fragment analysis

The Qubit Flex^TM^ Fluorometer model (ThermoFisher Scientific) with 1x dsDNA HS reagents was used to quantify DNA.

Fragment lengths from clinical samples and libraries were determined using the Fragment Analyser (Agilent) DNF-474-33 – HS NGS Fragment 1-6000bp or TapeStation (Agilent) High Sensitivity D1000 ScreenTape® or Genomic DNA ScreenTape®.

### Host DNA depletion

Host DNA depletion was performed on samples using the NEB Next® Microbiome DNA Enrichment Kit E2612L according to manufacturer’s instructions. For optimal performance, genomic DNA fragments should be 15 kb or longer. The DNA concentration was measured for each sample and, dependent on sample DNA concentrations, was added to the reaction in amounts ranging from 11.64 to 1000 ng, providing the maximum DNA input per sample. We normalised the sample concentrations to 1 ng/μL or 25 ng/μL according to the initial concentration, starting with an initial working volume of 20 μL. Samples containing the target microbial DNA were purified by AMPure XP beads (Beckman Coulter).

### SureSelect

Custom probes were designed around the reference genomes AM884177 (CT L2b/UCH-1), with or without LT591897 (NG WHOF), CP004010 (TP Nichols) and GCF_000027325_1 (MG G37). Further complete wild type genomes from Genbank (Table SX) were used to create probes to cover the known pan-genome of each species. Probes were designed with at least 90 % sequence homology to the reference genomes with 1-2X tiling frequency, by an inhouse developed algorithm by Ibrahim Kisakesen and Danielle Fletcher from Agilent Technologies (29,53). For regions with ambiguous bases, among all possible sequences, representative probes with 90% base homology threshold were selected. The algorithm also reduces the number of relevant baits to ensure cost efficiency. The final bait sets comprised 94874 baits (CT only, Design ID S3465202) and 242000 baits (CT, NG, MG, TP, Design ID S3465164).

Target enrichment was performed on clinical samples with SureSelect XT HS2 DNA with post-capture pooling for Illumina Platform NGS (G9983-90500 Rev A0, Agilent SureSelect). The protocol was performed as per the manufacturer’s instructions. For the first step of enzymatic DNA fragmentation, an initial volume of 17 µl was selected due to low DNA concentrations and manufacturers’ recommendations. The final volume was adapted to this modification to reach a total of 50 µL. With this modification the total DNA input ranged from 10 to 200 ng. For all thermal cycler programs, we used the T Professional TRIO Thermocycler (Biometra). The 37°C incubation step of the enzymatic fragmentation duration was performed for 15 minutes. The second stage, library preparation, used 11 PCR cycles for pre-hybridisation PCR. For the hybridization and capture step, to maximise the DNA input, we consistently used the maximum sample volume containing up to 1000 ng DNA made up to 12 µL with nuclease-free water for each sample. The probes were used undiluted or diluted 1:5 or 1:10. Hybridisation was performed overnight at 65°C. Post-hybridization used 16-22 PCR cycles.

### Illumina sequencing and bioinformatic analysis

Illumina sequencing was carried out PE150 on a NextSeq1000 platform after library generation with Qiaseq FX DNA library kit (Qiagen) or SureSelect (as above). Reads were trimmed with trimmomatic v0.39 (54). Mapping to relevant references (AM884177 (CT), LT591897 (NG), CP004010 (TP) and GCF_000027325_1 (MG), base calling and generating of multiple sequence alignment (MSA) used snippy v4.6.0 (https://github.com/tseemann/snippy) with default parameters except setting the minimum number of reads covering a site to be considered (mincov) to 5 to enable calling from regions of lower read depth. Whole genome sequencing was considered successful (“Genome success”) when percentage genome covered >95%; mean read depth >10x, and <20% Ns. On-target-read percentage (OTR%) was calculated using the fraction of mapped reads from the bam file generated by snippy using samtools v1.15.1 (55). Kraken2 v2.1.2 was used to determine the constituents of sequence data (56).

### ONT sequencing, adaptive sampling and bioinformatic analysis

For serial dilutions, cultured CT DNA from two HUCH and one UBA sample was spiked into human DNA (Promega G1471, diluted to 5 µg/µL for use as a working solution) and two serial dilutions were performed in human DNA.

Samples underwent rapid barcoding (Rapid Barcoding Kit 24 V14, SQK-RBK114.24) with double quantities and was halved to provide identical libraries for sequencing (9- to 10-plex) on the GridIon with and without adaptive sampling. Each R10.4 flow cell started with >1000 available pores. Adaptive sampling was used in the positive mode based on reference genomes AM884177 (CT L2b/UCH-1), with or without LT591897 (NG WHOF), CP004010 (TP Nichols) and GCF_000027325_1 (MG G37). Bases were called using the Dorado Super accuracy (dna_r10.4.1_e8.2_5khz_400bps_sup@v4.2.0) model, and samples demultiplexed with the inbuilt MinKnow software v23.04.5 in the GridION. Nanoplot v1.40.0 was used to access the raw read quality. Reads were selected to be >Q10 using nanofilt v2.8.0 (57). Mapping to the reference genomes as above used minimap2 v2.24 (58). %On-Target-Bases (%OTB) were calculated from minimap2 covbases and meandepth and nanoplot total bases >Q10.

### Figures and Statistical Analysis

Data was plotted using ggplot2 v3.4.4 (Wickham 2016) in R v4.3.2 (59) within RStudio 2023.09.1+494 (2023.09.1+494) (60). All statistical analyses were performed in R as above. To determine statistical differences between groups, Analysis of Variance (ANOVA) was used. Linear regression analysis was conducted to assess the relationship between pathogen load and genome success rate. p-values < 0.05 indicated that the relationship between Ct and %OTR is statistically significant and can be explained by a linear model. R^2^ was used to evaluate the fit of the regression model. Data normality was assessed using the Shapiro-Wilk test, and the non-parametric Mann-Whitney test was used to compare groups. p-values < 0.05 were considered statistically significant.

## Results and Discussion

### Metagenomic sequencing of clinical swabs samples

Clinical samples are often diagnosed using RT-PCR methods, which provide a cycle threshold (Ct) value as an indication of pathogen load. Therefore, from the diagnostic methods alone, it is known that the amounts of pathogen RT-PCR targets are very low (Table S2). The 14 Swiss clinical samples analysed were from mainly asymptomatic individuals, including seven samples positive for CT and four samples positive for NG, with Ct values over 25, three samples positive for MG with Ct values over 30, with only one TP positive sample with Ct of 29.9. In contrast, the 39 Argentinian clinical samples from symptomatic patients had higher loads of CT (39 positive samples) with 16 samples having Ct<25, and seven NG samples of which two had Ct<25. The four TP positive samples had Ct values from 27.2 to 36.4, and the two MG positive samples,had Ct values between 29.6-35.9.

In order to calculate the scale of the problem of generating genomes directly from these clinical samples, without culture, and to see whether metagenomic (deep) sequencing alone can provide genomic data on the target pathogens, we used 33 clinical samples from both Argentina and Switzerland, with a range of Ct values, and performed Illumina sequencing, with up to 31 million trimmed reads per sample. Of these, 30 were positive for CT; nine for NG; five for TP and three for MG. The resulting %On-Target-Reads (%OTR) shows that for samples diagnosed as positive for the four pathogens, %OTR were very low, ranging from 0.0001-0.012 for CT; 0.00004-0.25 for NG, 0-0.006 for TP and 0-0.003 for MG (Figure 1A). This compares with previous data on CT, being present in urine and vaginal swabs at <0.6% (61). A correlation was found between Ct values, estimating bacterial load, and %OTR in CT (p-value 4.9e-05), data was too limited for the other microorganisms studied (Figure 1B). The highest genome coverage for CT achieved by this method was 21% with a mean read depth of 0.3x; for NG 33% with mean read depth of 0.6x; and for other pathogens much lower, which is clearly insufficient for confident genomic analysis.

**Figure 1.**
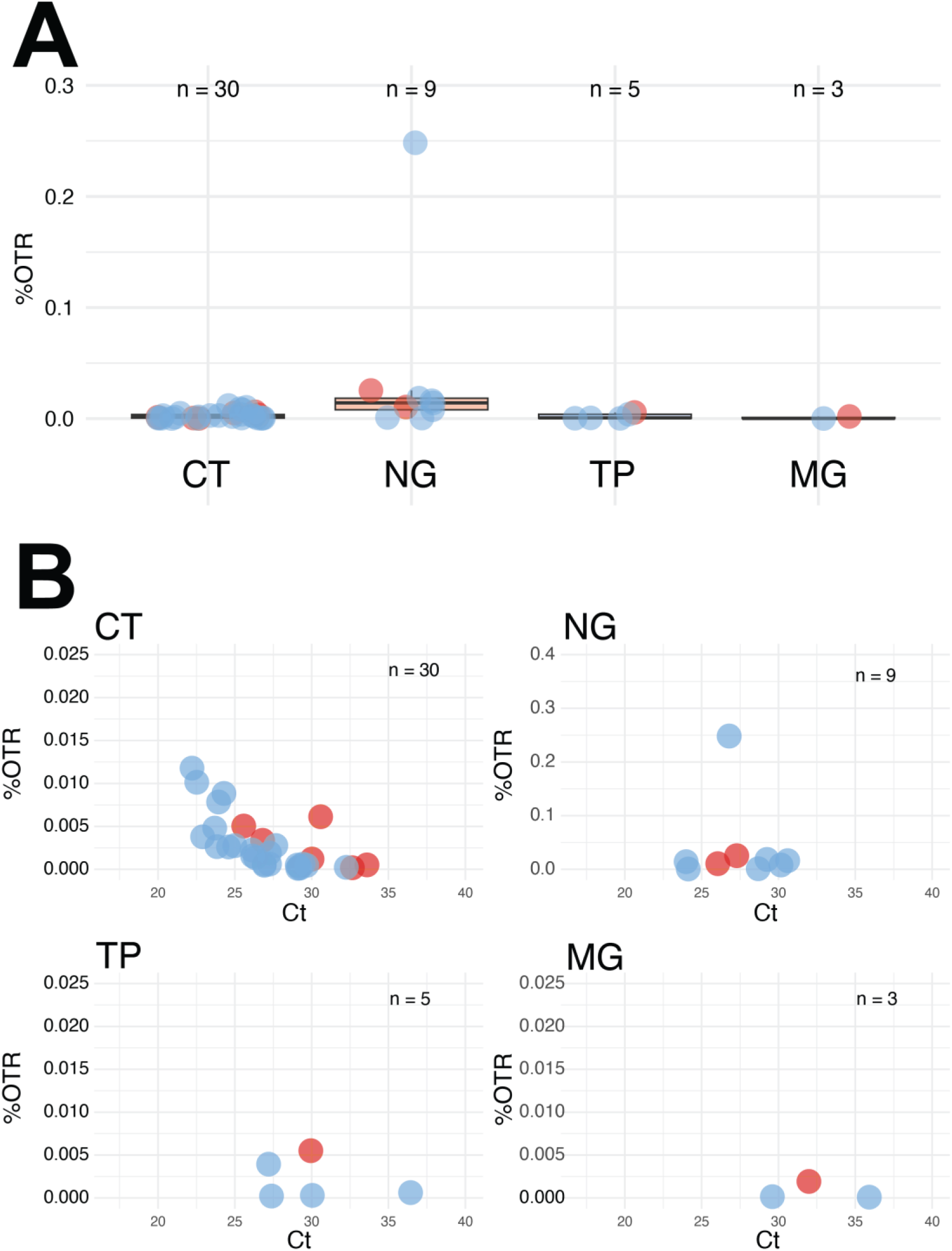
Direct metagenomic sequencing results. **A.** %OTR of each directly sequenced sample, for each pathogen (positively diagnosed samples). The line shows the median and the box 50% interquartile range (IQR). **B.** Correlation of %OTR with diagnostic RT-PCR Ct values for each positively diagnosed pathogen. Colours of data points reflect sample source: Argentina pale blue, Switzerland red. Please note the different scales on the y axis. With a p-value 4.9×10^-5^<< 0.05 the correlation between %OTR and Ct is statistically significant for CT.

### Host DNA depletion

Host DNA depletion, an untargeted approach which enriches all CpG-un-methylated DNA, was attempted on eight clinical samples (three from Argentina and five from Switzerland, all having undergone direct metagenomic sequencing) with the NEBNext® Microbiome DNA Enrichment Kit. This method is most effective on samples with DNA fragments over 15 kb. DNA fragment lengths were first assessed from these original extracted swab samples and three more (total six from Argentina and five from Switzerland). None of the Argentinian samples reached fragment lengths of 15 kb as required for the technique, while Swiss samples all had mean fragment lengths above those required (Table S3).

Of the eight clinical samples tested, all were diagnostically positive for CT and one also for NG. At these low pathogen loads, reads assigned to the species of interest (%OTR) did not exceed 0.015% in any sample (Figure 2). Thus this method of host DNA depletion did not provide better results than direct metagenomic sequencing of the sample. It cannot be determined from this data whether fragment lengths had an impact on the results.

**Figure 2.**
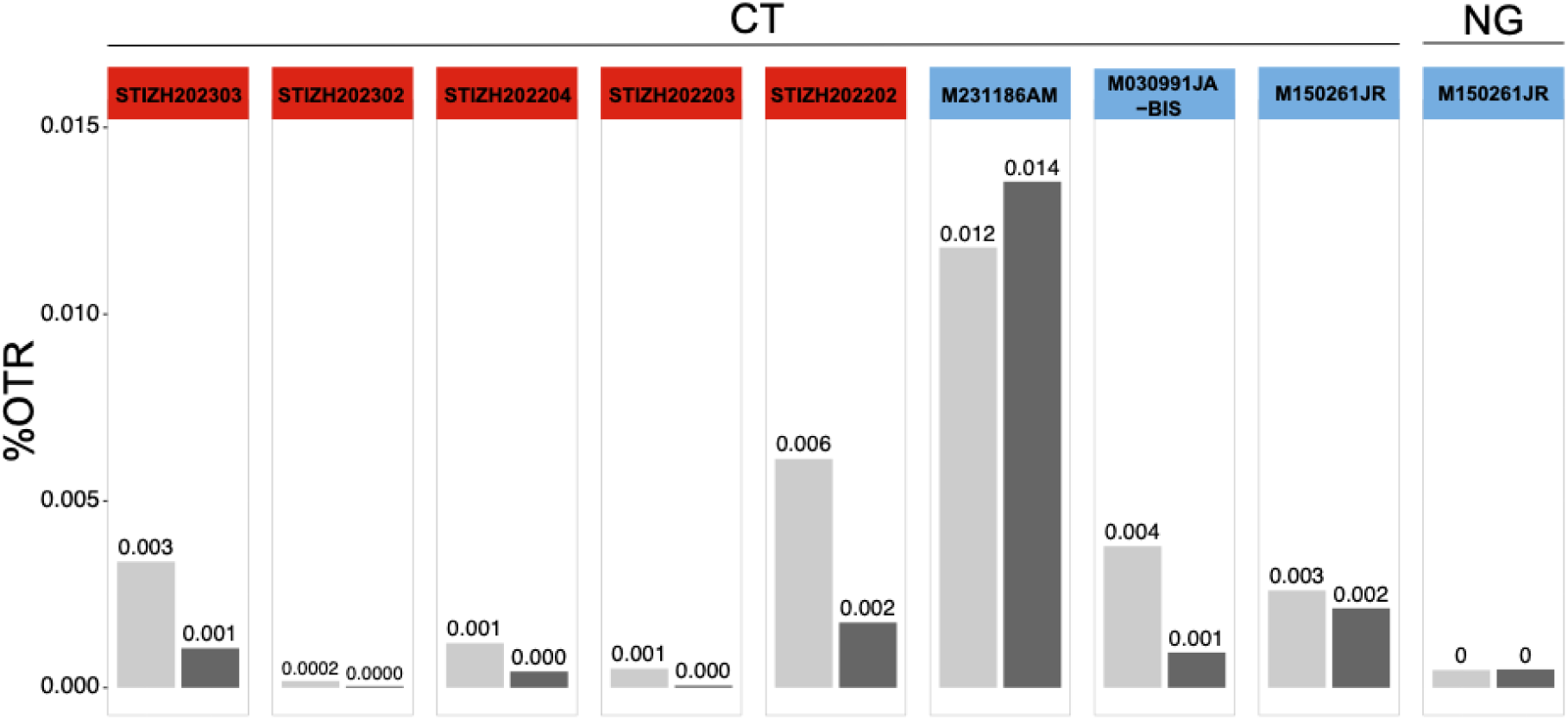
Comparison of host DNA depletion to direct metagenomic sequencing. %OTR for individual samples and pathogens (CT and NG). Pale grey bars show results of direct metagenomic sequencing, and dark grey bars sequencing after host DNA depletion. Colours of headers reflect sample source: Argentina pale blue, Switzerland red.

In addition to the sought STI pathogens, these anorectal samples will contain a considerable amount of microbial DNA as well as host DNA. In order to see if this was a factor in the lack of enrichment, we defined the reads assigned as human or bacterial in both metagenomic sequenced and host DNA depleted samples. In all samples except one, which contained more reads assigned to faecal genera, human reads made up over 94% and increased to over 97% after host DNA depletion and reads assigned to bacteria made up under 5% and decreased to under 1.1% after host DNA depletion (Table S4).

Given these results, this method is not suitable for generating genomes of STI bacteria from clinical samples. In extracted samples from Argentina and Switzerland issues likely to affect the success of the method were identified with DNA fragment length and concentration respectively. These differences are likely due to sample collection, processing and transport issues, including freeze-thawing cycles. The condition of the sample may also be partly responsible for the lack of success, but is also not possible to influence. In addition, the amount of human DNA present may have been overwhelming for this method.

### ONT adaptive sampling

A technology with much promise is the adaptive sequencing method from ONT, where only selected reads are fully sequenced, after mapping of a short region of the read against a supplied reference genome. This is therefore a targeted approach, which requires high quality reference genomes of the species in question.

Ten Swiss clinical samples were sequenced in parallel with and without adaptive sampling, selecting for reads using reference genomes from CT, NG, TP, and MG. Swiss samples were selected due to the longer fragment lengths. Five samples were positive for CT, one for NG, one for CT and NG, one for TP, and two were positive for MG. With or without adaptive enrichment, %On-Target-Bases (%OTB) was always under 0.1% (Table S5). Mean read lengths between were 1610 ± 638 without adaptive sampling and 575 ± 40 with adaptive sampling, many samples saw no enrichment, and the highest degree of enrichment observed was 2.1x. No data coming close to forming a whole genome was generated.

In a spike experiment, the method was found to work better on higher load samples. We added CT DNA from three cultured samples (two from HUCH, and one from UBA) into carrier human DNA at 4-54% CT (Table S6), from which two serial dilutions were made to cover some of the range of possible target concentrations in extracted clinical samples. Adaptive sampling was performed, selecting only for reads mapping to the CT reference. We obtained shorter mean read lengths with adaptive sequencing, as the shorter reads cannot be selected, and are fully sequenced whereas the longer reads are rejected (Table S6). CT reads were enriched to a maximum 66.7-fold in %OTB, but this did not improve the generation of genomes. When the data was filtered for longer (>2kb) reads which were analysed, fold-enrichment was seen to increase, but as a smaller portion of the data was used in this analysis, no genomes could be generated here either (data not shown).

This technology requires longer reads and higher target loads than available in our clinical samples for successful adaptive sampling. As many of the reads we generated were shorter than 500bp, and from extremely low pathogen load samples, we saw almost no enrichment effect. Unfortunately, given the condition and load within clinical samples, adaptive sampling is not useful for genome generation in this context. Previous studies on comparable, but non-STI material also showed low levels of enrichment, and failure to generate full genomes (62,63).

### Target enrichment of clinical samples with SureSelect

To date no data on target enrichment against a panel of STI-causing bacteria has been published. We designed a first iteration of target enrichment baits targeting only CT. A second panel of baits targeted all four pathogens CT, NG, TP, and MG and we attempted to obtain the genomes of four STI bacteria in one experiment.

We first performed CT-only enrichment, and then compared this to the panel bait enrichment, to see whether the panel causes a reduction in success rate. Here we had 13 clinical samples to compare in 29 experiments (Figure S1). The results between both bait sets are equivalent. For five samples (in13 replicate experiments), genomes were not obtained with either bait set. For six samples (in12 replicate experiments), whole genomes (see Methods for criteria) were obtained using both methods. For two samples (in four replicate experiments), whole genomes were produced using one set of baits, with read depth for the other bait set falling just under our acceptance criteria (Table S7). With higher sequencing coverage, the results would likely have been equivalent. Results across duplicate experiments were consistent, also in terms of SNP detection (64). Thus, the panel baits do not cause a reduction in success in capturing CT genomes.

**Figure S1.**
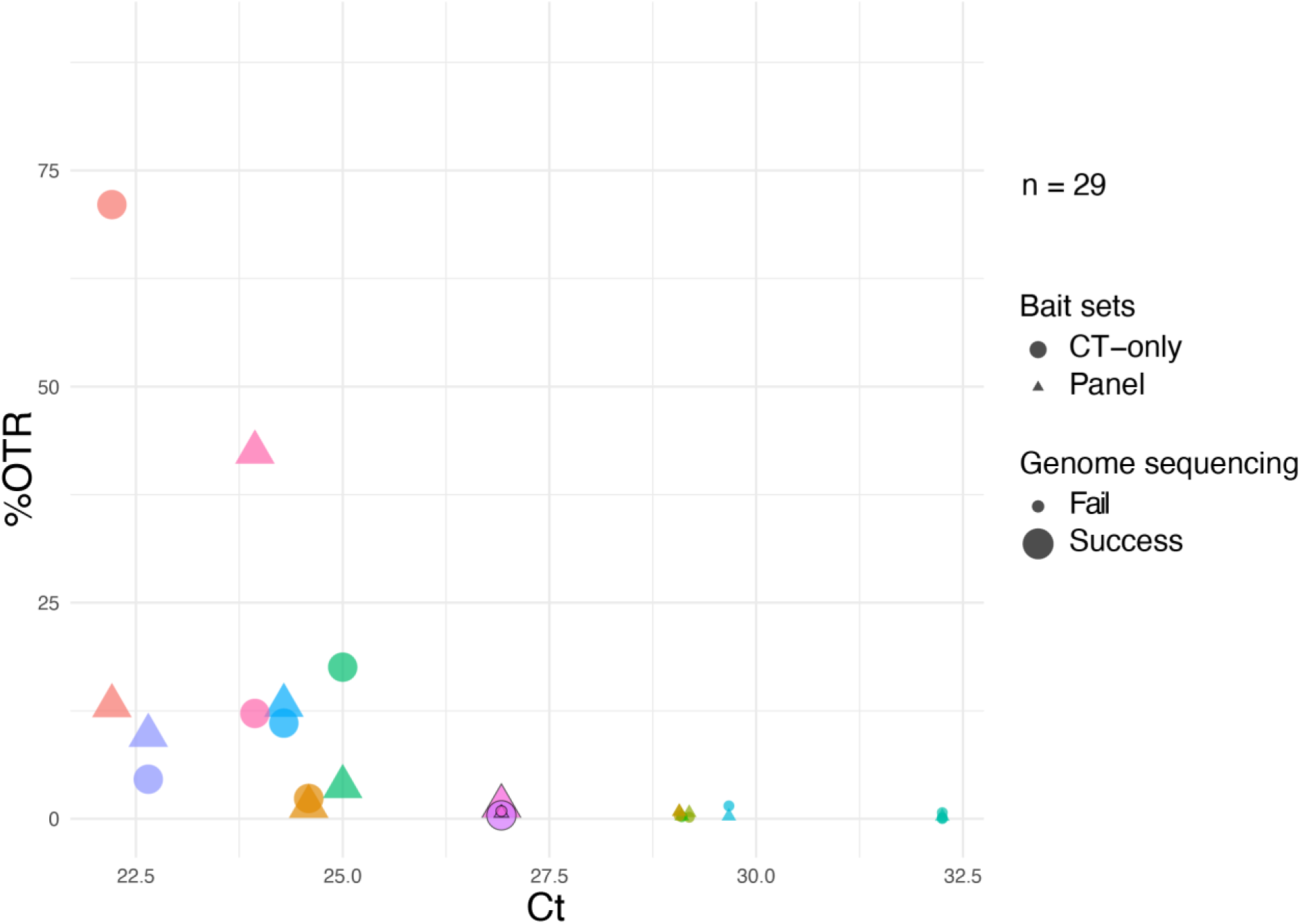
Comparison of CT data enrichment between CT-only and panel bait sets. Samples (13) over 29 experiments are labelled by colour. The two samples with different results for the baits sets are shown with black outlines. Genome sequencing success is defined as coverage >95% and mean read depth >10 (see Methods).

Our overall analysis looked at results from both CT-only and the panel baits, investigating the bacterial pathogen genomes that were expected to be present from diagnostic RT-PCR results. We investigated enrichment levels compared to direct metagenomic sequencing of clinical samples, and genome sequencing success rates, compared to bacterial load.

Comparing SureSelect to direct metagenomic sequencing, we used 32 samples of which 29 were positive for CT, nine for NG, five for TP, and three for MG. (Figure 3). SureSelect resulted in enrichment for each pathogen, and successful genomes were only obtained from the SureSelect method.

**Figure 3.**
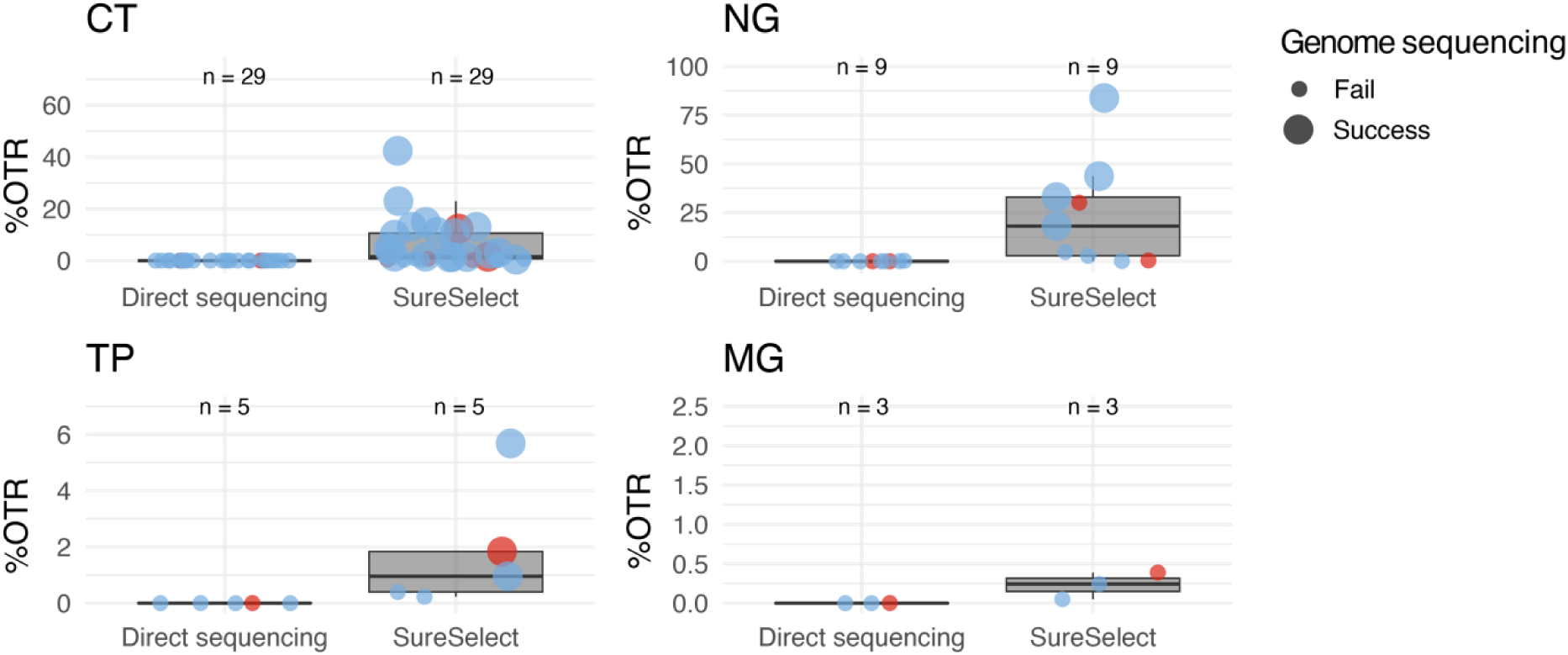
Enrichment of genomes using SureSelect compared to metagenomic sequencing. %OTR are used to show the degree of enrichment using SureSelect. Where replicate experiments were performed, the first panel bait experiment was used as a representative, except for samples M260278JB and M281090JR, where the successful samples with higher reads depth are represented. The line shows the median and the box 50% interquartile range (IQR). Colours of data points reflect sample source: Argentina pale blue, Switzerland red. Genome sequencing success, defined as coverage >95% and mean read depth >10 (see Methods), is represented by large points and failure by small points. Please note the different scales on the y axis.

To investigate genome success rates after SureSelect target enrichment, we were able to investigate 45 CT positive samples (17 with replicates), 11 NG positive samples (one duplicate), five TP positive samples and five MG positive samples (Figure 4). The success rates for CT were 64%, for NG 36% (with a further two samples providing over 90% coverage), for TP 60% and for MG 0%. Replicate experiments showed overwhelmingly the same results, with the exception of two CT samples, where increased sequencing depth allowed successful genome generation. Three MG samples (60%) showed >65% coverage and >5x mean read depth; given the extremely low %G+C in this genome a reduced temperature during the capture protocol could improve these results.

**Figure 4.**
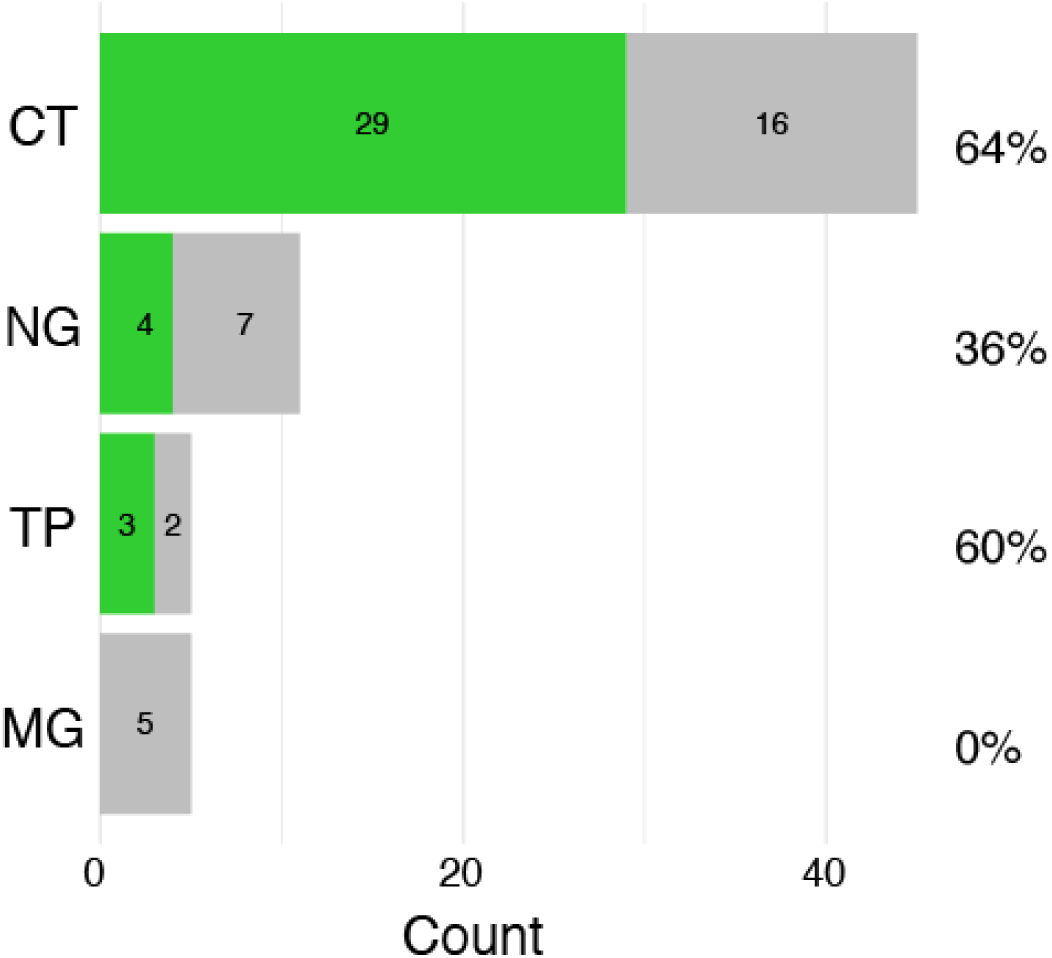
Success of genome enrichment using SureSelect for the four pathogens CT, NG, TP, and MG. Successfully genome sequenced samples are shown in green and unsuccessful samples in grey. Genome sequencing success is defined as coverage >95% and mean read depth >10.

For the above samples, we compared %OTR after target enrichment to the diagnostic RT-PCR Ct values, correlated with the defined success of genome sequencing. For CT, where we have most data, we observe a significant correlation between pathogen load and success rate (p << 0.05; R^2^ 0.45, x intercept 29.4), suggesting a Ct value cutoff for success around Ct30 (Figure 5), comparable to that described in previous publications (22,27). Replicate experiments of the same samples may show different %OTR data but the genome sequencing success or failure remains the same (data not shown).

**Figure 5.**
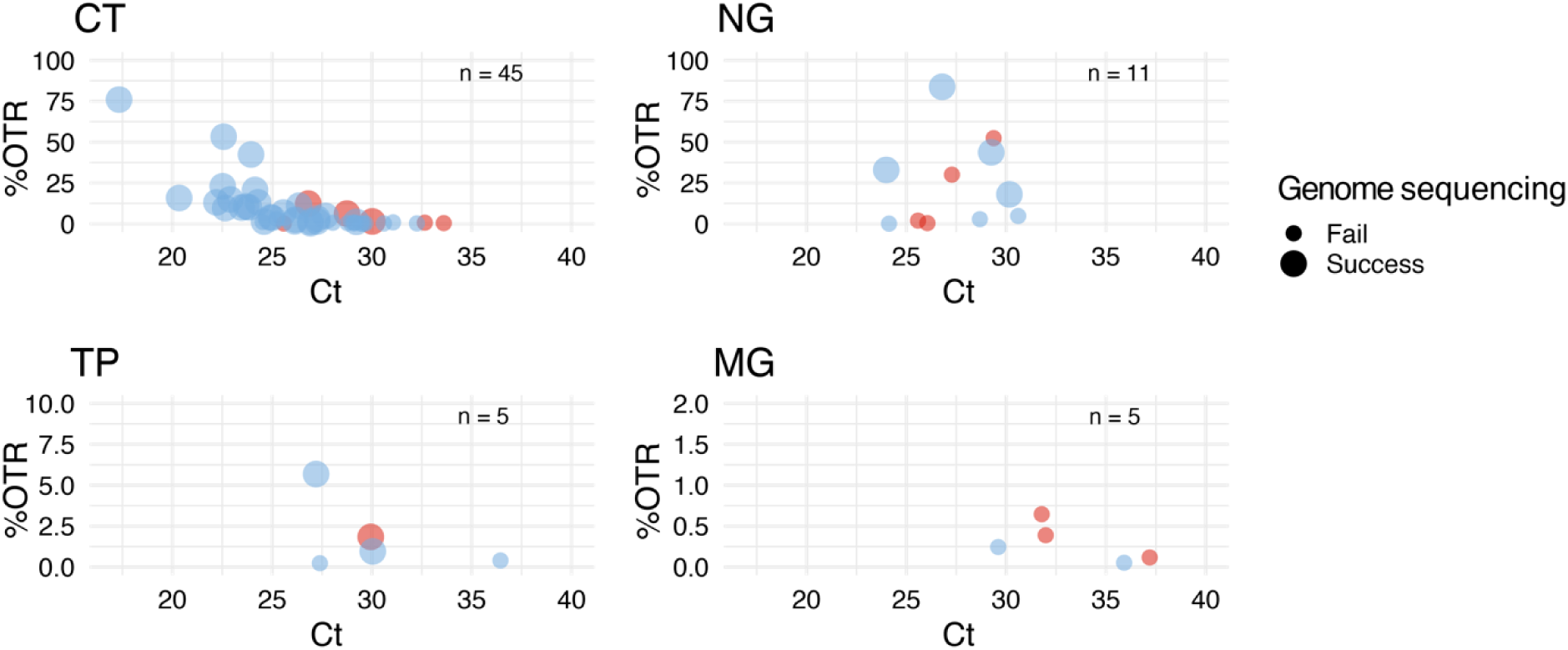
Correlation of %OTR from SureSelect with diagnostic RT-PCR Ct values. Colours of data points reflect sample source: Argentina pale blue, Switzerland red. Genome sequencing success, defined as coverage >95% and mean read depth >10 (see Methods), is represented by large points and failure by small points. Please note the different scales on the y axis. With a p-value 4.4×10^-7^<< 0.05 the correlation between %OTR and Ct is statistically significant and can be explained by a linear model with R^2^ of 0.45 and an x intercept of 29.4. The p-values for other species were limited by lack of data and were not significant.

For the other three pathogens, we have less data, and correlations are not as clear, but the trend of genome success below around Ct30 also holds (Figure 5).

The cost of target enrichment using custom designed probes is indeed relatively high, but offers results that are not possible with other techniques. Our method used post-capture pooling to optimise the chances of obtaining whole genomes, especially from co-infected samples, with bait dilution to mitigate the costs of this reagent. Cost calculations and comparisons are complicated, and experimental success needs to be factored in (65).

## Conclusions

We present here a panel bait approach, where the genomes of four major STI bacterial pathogens from clinical samples can be targeted in a single experiment. Enrichment of bacterial STI genomes in our hands was effective only using target enrichment. Deep metagenomic sequencing, host DNA depletion, and ONT adaptive sampling did not produce more than 0.25% OTR. A previous publication describes the generation of five(98% complete *C. trachomatis* genomes from 138 metagenomic sequencing experiments (66), which would not be possible at the pathogen loads in our samples. We obtained full genomes only using SureSelect target enrichment (CT from 64% of clinical samples, NG 36% and TP 60%). We also intend to optimise our experimental procedure to increase the capture of MG genomes.

Given the collection, storage, and transport variables of clinical samples, an advantage of target enrichment compared to two of the alternative techniques described here is that there is no requirement for high molecular weight DNA. Even short fragments can be used to generate libraries and undergo capture. The samples from Argentina, being from symptomatic patients in comparison to the Swiss screening samples, tended to have higher load (lower Ct values), which may increase the success of the enrichment. Another factor aiding successful genome generation in borderline samples can be the read depth: for two samples we found that increased read depth was the difference between passing and failing our criteria for genome completeness. It has also been described that SureSelect can pick up in-host variation within genomes (67). With careful bioinformatic analysis, this can be an advantage, but this was not seen in our genomes.

The success of this approach means that this technique can be used by other laboratories and on further global samples to inform on circulating STI strains without the need for unreliable and labour intensive culture (64). Rapid knowledge on the AMR determinants of detected STI-pathogens would also allow a more targeted approach regarding treatment decision and also for antibiotic stewardship. Subsequent analysis of the genomes in the context of previously published genomes will also provide important insights. This novel STI-panel genome target enrichment approach can provide epidemiological data directly from the sample.

## Supplementary files

Table S1. List of genome plasmid accessions used for probe design.

Table S2. Description of samples included in the study.

Table S3. Fragment lengths of 11 clinical samples.

Table S4. Kraken2 results of paired samples analysed by direct sequencing (-ns) or host DNA depletion (-HD).

Table S5. Results of ONT adaptive sampling experiment on clinical samples.

Table S6. Results of ONT adaptive sampling experiment on CT spiked samples.

Table S7. Comparison of CT baits against panel baits for 13 samples.

## Author contributions

KB: Investigation, Data curation, Formal Analysis, Methodology, Visualization

VB: Investigation, Formal Analysis, Methodology, Visualization

FW: Software, Formal Analysis, Visualization

SP: Investigation

FI: Resources, Investigation

TR: Data curation

MHP: Resources

EHB: Resources

DB: Resources

IK: Methodology

AS: Methodology

ACE: Resources

MLGV: Resources

DLA: Resources

LSL: Resources

LLR: Resources

AE: Resources, Writing – review & editing

MRF: Funding acquisition, Project administration, Supervision

HMBSS: Conceptualization, Data curation, Formal Analysis, Funding acquisition, Methodology, Project administration, Supervision, Validation, Writing – original draft

## Ethical approval

No patient data is mentioned in this paper and all samples are deidentified. As such, no specific ethical approval is required.

## Consent for publication

No consent for publication was required.

## Acknowledgements

We thank Daniel Gander, Valéria Pires, Arianita Asani, and Stefan Antener for excellent technical assistance with sequencing. TapeStation data produced and analyzed in this paper were generated in collaboration with the Genetic Diversity Centre (GDC), ETH Zurich. Many thanks to Sofia Polcowñuk for designing the graphical abstract.

